# Demystifying image-based machine learning: A practical guide to automated analysis of field imagery using modern machine learning tools

**DOI:** 10.1101/2022.12.24.521836

**Authors:** Byron T. Belcher, Eliana H. Bower, Benjamin Burford, Maria Rosa Celis, Ashkaan K. Fahimipour, Isabella L. Guevara, Kakani Katija, Zulekha Khokhar, Anjana Manjunath, Samuel Nelson, Simone Olivetti, Eric Orenstein, Mohamad H. Saleh, Brayan Vaca, Salma Valladares, Stella A. Hein, Andrew M. Hein

## Abstract

Image-based machine learning methods are quickly becoming among the most widely-used forms of data analysis across science, technology, and engineering. These methods are powerful because they can rapidly and automatically extract rich contextual and spatial information from images, a process that has historically required a large amount of manual labor. The potential of image-based machine learning methods to change how researchers study the ocean has been demonstrated through a diverse range of recent applications. However, despite their promise, machine learning tools are still under-exploited in many domains including species and environmental monitoring, biodiversity surveys, fisheries abundance and size estimation, rare event and species detection, the study of wild animal behavior, and citizen science. Our objective in this article is to provide an approachable, application-oriented guide to help researchers apply image-based machine learning methods effectively to their own research problems. Using a case study, we describe how to prepare data, train and deploy models, and avoid common pitfalls that can cause models to underperform. Importantly, we discuss how to diagnose problems that can cause poor model performance on new imagery to build robust tools that can vastly accelerate data acquisition in the marine realm. Code to perform our analyses is provided at https://github.com/heinsense2/AIO_CaseStudy

## Introduction

Imagery from the ocean has long been a tool for surveying marine environments, quantifying physical conditions, and monitoring the inhabitants of marine ecosystems (Longley & Martin 1927, Drew 1977, Beijbom *et al*. 2015, Lombard *et al*. 2019, Marochov *et al*. 2021). This reliance on imagery as a means of extracting data from marine systems has only grown with the increasing accessibility of satellite imagery and the decreasing cost and increasing quality of imaging systems that can be deployed directly in the field, including ultra-low cost “action cameras” (Williams *et al*. 2019, Rodriguez-Ramirez *et al*. 2020, Bamford *et al*. 2020, Durden *et al*. 2016). Yet visual data bring with them some unique challenges. Images and video are expensive to process due in part to the fact that imagery is inherently high-dimensional; for example, a single grayscale image of one-megapixel resolution, a coarse image by modern standards, is a 2^20^-dimensional data object. Researchers who collect images as their source of raw data often return from field campaigns with terabytes or even petabytes of image data that need to be processed.

The role of ***image analysis*** (see Table 1 for glossary of bolded terms) is to compress high-dimensional visual data into much lower-dimensional summaries relevant to a particular task or study objective. For example, if the goal of a study is to determine whether a particular species of interest is or is not present, all of the relevant data in an image can be compressed to a single bit of information that encodes whether the species is or is not present in the image. As humans, we perform this type of visual data compression naturally (Marr 1982). We look at an image and with proper training, can classify what is present in the image, localize and count distinct objects, and partition the image into regions of one type or another. The objective of image-based ***machine learning*** (ML) is to train computer algorithms to perform these same tasks with a high level of accuracy. Doing so can tremendously accelerate image processing and greatly reduce its cost (Norouzzadeh *et al*. 2018), while also providing an explicit, standardized, and reproducible workflow that can be shared easily among researchers and applied to new problems (Goodwin *et al*. 2022, Katija *et al*. 2022). Despite the promise of these methods, the expertise required to apply, adapt, and troubleshoot ML methods using the kinds of image datasets marine scientists collect still represents a high barrier to entry.

**Table 1.** Glossary of terms relevant to image-based machine learning.

| **Term** | **Definition** |
| --- | --- |
| ***Image analysis*** | The process of extracting task-relevant information from imagery. |
| ***Machine learning*** | A body of mathematical and computational methods for extracting information from data to make predictions. |
| ***Machine learning pipeline*** | A computer program or set of programs that reads in training data, specifies and trains an ML model, produces model predictions, and provides performance metrics. |
| ***Image annotation*** | Process of generating ground truth labels for images, which are typically used to train ML models or evaluate performance. |
| ***Ground truth*** | A verified record, often produced by a human annotator, that describes what is contained within the image. Sometimes also called an annotation, or label. |
| ***Image classification*** | A task in which a whole image is assigned a class from a list of valid classes. |
| ***Object detection*** | A task in which objects within a set of classes of interest are detected and localized within an image, typically either within a bounding box, or polygon region. Many object detection methods also classify objects. |
| ***Instance segmentation*** | A task in which individual instances of objects in a class or classes of interest are localized within an image. Sometimes used synonymously with object detection, when objects are localized within polygons rather than bounding boxes. |
| ***Semantic segmentation*** | A task in which all individual pixels in an image are assigned to a class, but individual instances of objects are not specified. |
| ***Supervised learning*** | A type of machine learning that involves training a model with example input-output pairs. |
| ***Panoptic labels*** | A type of annotation that assigns a class to each pixel in an image and delineates the borders of instances of distinct objects of interest. |
| ***Few-shot learning*** | Machine learning methods designed to achieve good performance by training on few examples. |
| ***Deep neural network (DNN)*** | A machine learning method based on networks of interconnected computing nodes called "neurons." DNNs take data as input, process the data through one or more sequential layers of processing known as "hidden layers," and return predictions about the image. |
| ***Classification accuracy*** | Fraction of class predictions that are correct: (true positives + true negatives) / total number of predictions. |
| ***Precision*** | The fraction of positive class predictions that are correct: true positives / total predicted positives. |
| ***Recall*** | The fraction of positives present in the dataset that are correctly predicted by a model: true positives / total positives present in dataset. Sometimes referred to as *sensitivity*. |
| ***F1 score*** | A performance measure that incorporates both precision and recall: $2 \text{ (precision} \times \text{recall)} / (\text{precision} + \text{recall})$. |
| ***Intersection-over-union (IoU)*** | A measure of spatial localization performance used in object detection and instance segmentation. IoU measures the number of pixels contained within both the predicted instance location and the ground truth ("intersection" between the two areas), divided by the total number of unique pixels contained within the predicted instance location, and ground truth ("union" of the two areas). |
| ***mean average precision (mAP)*** | An average measure of classifier performance when bounding boxes or object instances are classified. Incorporates precision, recall, and IoU. |
| ***k-fold cross validation*** | A type of model evaluation in which training, validation, and test data are partitioned into $k$ different splits, and performance measures are evaluated on each split. |
| ***Distribution shift*** | Systematic differences in image statistics, scene complexity, class identities and distributions, and other relevant features between a training set and a new dataset to which a model is to be applied. |
| ***Image augmentation*** | The process of applying random digital alterations to training imagery during the training process to improve model generalization. |
| ***Image resolution*** | The resolution of the image in pixels. Many ML pipelines reduce image resolution by default to save memory and reduce training and deployment times. |
| ***Background imagery*** | Images that do not contain classes of interest. |
| ***Class coarsening*** | The process of lowering the resolution of classes by grouping several fine classes (e.g., species A, B, C, and D) into coarser classes (e.g., genus 1, genus 2). |

A number of review articles published in the last several years provide overviews of how modern image-based machine learning methods work and how these methods have been applied to problems in marine science (*e*.*g*., Goodwin *et al. 2022*, Li *et al*. 2022, Michaels *et al*. 2019). Here, we focus on the problem of practical implementation of image-based ML pipelines on real imagery from the field. The remainder of this paper is structured as a sequence of steps involved in defining an analytical task to be solved, preparing training data, training and evaluating models, deploying them on new data, and diagnosing performance issues. To provide concreteness, we present a running case study: object detection of marine species using imagery and software tools from the open source *FathomNet* database and interface (Katija *et al*. 2022). We use this case study to demonstrate each phase of constructing and troubleshooting an ML pipeline, and we provide code and guidelines needed to reproduce each step in the form of Google Collaboratory (Colab) notebooks (https://github.com/heinsense2/AIO_CaseStudy). We also provide performance benchmarks of this code on local hardware, and on free cloud computing resources (Google Colab cloud). Cloud computing resources enable those who wish to begin using these methods immediately to do so without the need to purchase specialized computing hardware.

## Training and using a machine learning pipeline

Researchers often have a clear idea of how they want to use the data extracted from imagery, and this idea forms the starting point for designing a ***machine learning pipeline*** to automatically extract data from imagery. Building a machine learning pipeline to adaptively to solve image analysis tasks involves a series of steps:

1. **Define an analytical task**. This step involves working to carefully define the objective of image analysis and the target metrics to be extracted from imagery. The type of imagery to be analyzed should be specified. This step may also involve defining performance criteria and setting benchmarks for acceptable performance.
2. **Generate and curate training and testing datasets**. This step involves developing and organizing image libraries for training, testing, and deploying models. This involves both organizing imagery with appropriate file structure and, very often, hand-labeling ***ground truth*** data to be used to train and test models. This step requires software tools that allow a researcher to organize images and to label, or ***annotate***, imagery so it can be later used to train and test machine learning models.
3. **Select and train appropriate machine learning models**. This step requires identifying a machine learning model architecture capable of performing the desired image analysis task. This step also involves identifying software and hardware implementations capable of training and deploying the model to perform inference on new imagery. When training, data must be partitioned into training, validation, and testing sets, and software tools may be needed to perform this partitioning in a customizable way.
4. **Evaluate model performance**. This step involves summarizing and visualizing model predictions and performance measures, and often comparing these measures across alternative model architectures or training schedules.
5. **Diagnose performance issues and apply interventions to improve performance**. This step involves applying a trained model to new imagery and re-evaluating its performance. If performance is below target levels, it may be necessary to modify training methods, datasets, or model architecture to improve performance.

In the following sections, we walk through each of these steps to illustrate how each is accomplished, and how the steps combine to produce an adaptable pipeline with robust performance.

## (1) Defining the image analysis task

### 1.1 Overview

Defining the image analysis task to be solved is the first step in any machine learning pipeline. Is the goal to assign an image to one class or another – for example, to decide whether a particular species is or is not present or a particular environmental condition is or is not met? Or is the aim instead to identify and count objects of interest – for example, to find all crustaceans in an image and identify them to genus? Or is the objective to divide regions of the image into distinct types and quantify the prevalence of those types – for example, to partition the fraction of a benthic image occupied by different algae or coral morphotypes (Dumas *et al*. 2009)? The answers to these questions determine how one proceeds with gathering appropriate labeled data, selecting and training a model, and deploying that model on new data.

### 1.2 Technical considerations

Many of the traditional problems marine scientists currently use imagery to address fall into one of three categories: *image classification, object detection, or semantic segmentation*. More complex tasks such as tracking (Irisson *et al*. 2022, Katija *et al*. 2021), functional trait analysis (Orenstein *et al*. 2022), pose estimation (Mathis *et al*. 2020), and automated measurements (Fernandes *et al*. 2020) often rely on these more basic tasks as building blocks. In what follows, we will focus primarily on these three core tasks, and reference other applications of image-based ML where appropriate.

In ***image classification*** problems, a computer program is presented with an image and asked to assign the image to one of a set of classes. Classes could be defined based on the presence or absence of particular objects (e.g., shark present or shark absent; Sharma *et al*. 2018), or represent a potentially broad set of categories to which the image must be assigned. An important distinction between whole-image classification and other common image analysis tasks is that in image classification, classes are assigned at the scale of the entire image (Fei-Fei *et al*. 2004, Chapelle *et al*. 1999). Thus, objects of interest are not spatially localized within the image, and the model does not provide information on the properties of individual pixels or spatial regions within the image. Whole image classification is appropriate for some tasks, such as simply detecting the presence or absence of a particular species of interest or environmental condition, but is less appropriate for others, for example, counting individuals of a particular species when multiple individuals can occur within a single image (Beery *et al*. 2021). Cases in which multiple object classes can occur in the same image can also pose problems for whole image classification tasks. Nevertheless, this task remains relevant in automated image analysis problems (Qin *et al*. 2016, Villon *et al*. 2021, Kyathanahally *et al*. 2022) and is the approach of choice for certain types of marine microscopy data where the images are typically stored as extracted region of interest (e.g., Luo *et al*. 2018, Ellen *et al*. 2019).

A second common task involves detecting and spatially localizing objects of interest within images, a task known as ***object detection*** or ***instance segmentation***. Separating instances of the same type of object (e.g., there are nine fish identified as Atlantic cod in this image) in a given image is often crucial if imagery is being used to estimate abundances (Moeller *et al*. 2018, but see Scoulding *et al*. 2022 for a discussion of limitations at high density), and most object detection pipelines can be trained to detect objects of many different classes, which is valuable for analyzing images that contain multiple objects of interest of different types (e.g., many distinct phytoplankton species or morphotypes in the same image, Irisson *et al*. 2022).

A third task that is relevant for many marine science applications involves assigning a class to each pixel in an image, a task known as ***semantic segmentation***. Semantic segmentation differs from object detection in that, under semantic segmentation, one is not interested in detecting and discriminating instances of a particular class, but rather determining the class membership of each pixel in an image. This can be useful for tasks such as estimating the percent cover of algae, corals, or other benthic substrate types (e.g., Beijbom et al. 2015, Williams *et al*. 2019). If images are collected in a controlled and standardized way, the percentage of each image occupied by different species or classes of object can be estimated by the relative abundance of pixels assigned to each class.

Image-based ML tools have also been used for a variety of applications beyond the three tasks described above. Examples include “structure-from-motion” studies, in which the three-dimensional structure of objects are inferred and reconstructed from a sequence of images taken from different locations in the environment (Francisco *et al*. 2020), animal tracking and visual field reconstruction (Hein *et al*. 2018, Fahimipour *et al*. 2022), quantitative measurement and size estimation (Fernandes *et al*. 2020), animal postural analysis (Graving *et al*. 2019), and re-identification of individual animals in new images based on a set of previous observations (Nepovinnykh *et al*. 2020).

### 1.3. Case study: species detection and classification from benthic and midwater imagery

To provide a concrete example, throughout the rest of this review, we will consider an object detection and classification task that seeks to localize and identify marine animals in deep-sea imagery collected from the Eastern Pacific, within the Monterey Bay, and surrounding regions. Images were collected by the Monterey Bay Aquarium Research Institute (MBARI) during Remotely Operated Vehicle (ROV) surveys conducted between 1989 and 2021 (Robison *et al*. 2017), and are housed in the open-source *FathomNet* database (FathomNet.org; Katija *et al*. 2022). We focus on six common biological taxa that are observed broadly across the sampling domain, at a range of depths, and over several decades of sampling (Fig. 1. shows iconic image of each class): the fish genera *Sebastes* (Rockfish) and *Sebastolobus* (Thornyheads), and the squid species *Dosidicus gigas* (Humboldt squid), *Chiroteuthis calyx* (swordtail squid), *Gonatus onyx* (black-eyed squid), and the siphonophore, *Nanomia bijuga*. Although classes of interest are sometimes clearly visible in images as shown in Figure 1, *FathomNet* contains many images with small subjects, complex visual backgrounds, heterogeneous lighting, and a host of other challenging visual conditions (Figure 2). Importantly, these conditions are characteristic of the imagery collected in many marine science applications, and they make reliable detection, localization, and classification of objects of interest a non-trivial task.

**Figure 1.**
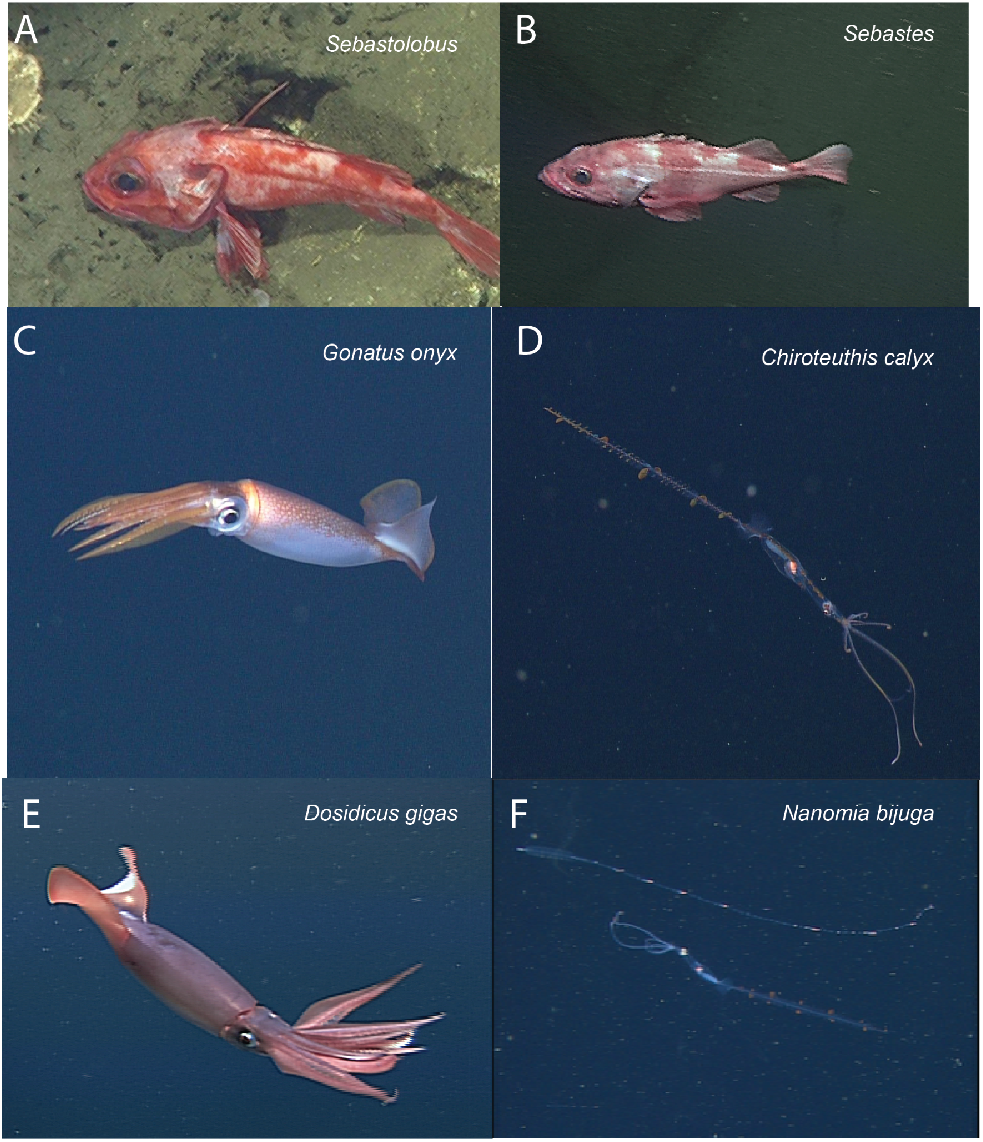
Focal species included in case study (iconic images). Focal species included fish in the genera *Sebastolobus* (A) and *Sebastes* (B), squid species *Gonatus onyx* (C) *Chiroteuthis calyx* (D), and *Dosidicus gigas* (E). Panel (F) shows an image of the siphonophore, *Nanomia bijuga*, alongside a juvenile *C. calyx* (F, lower organism in image), which are believed to visually and behaviorally mimic *N. bijuga* (Burford *et al*. 2015). Images in panels A-F were selected for clarity and subjects are enlarged for visualization. Figure 2 shows focal species in images that are more representative of typical images in *FathomNet*.

**Figure 2.**
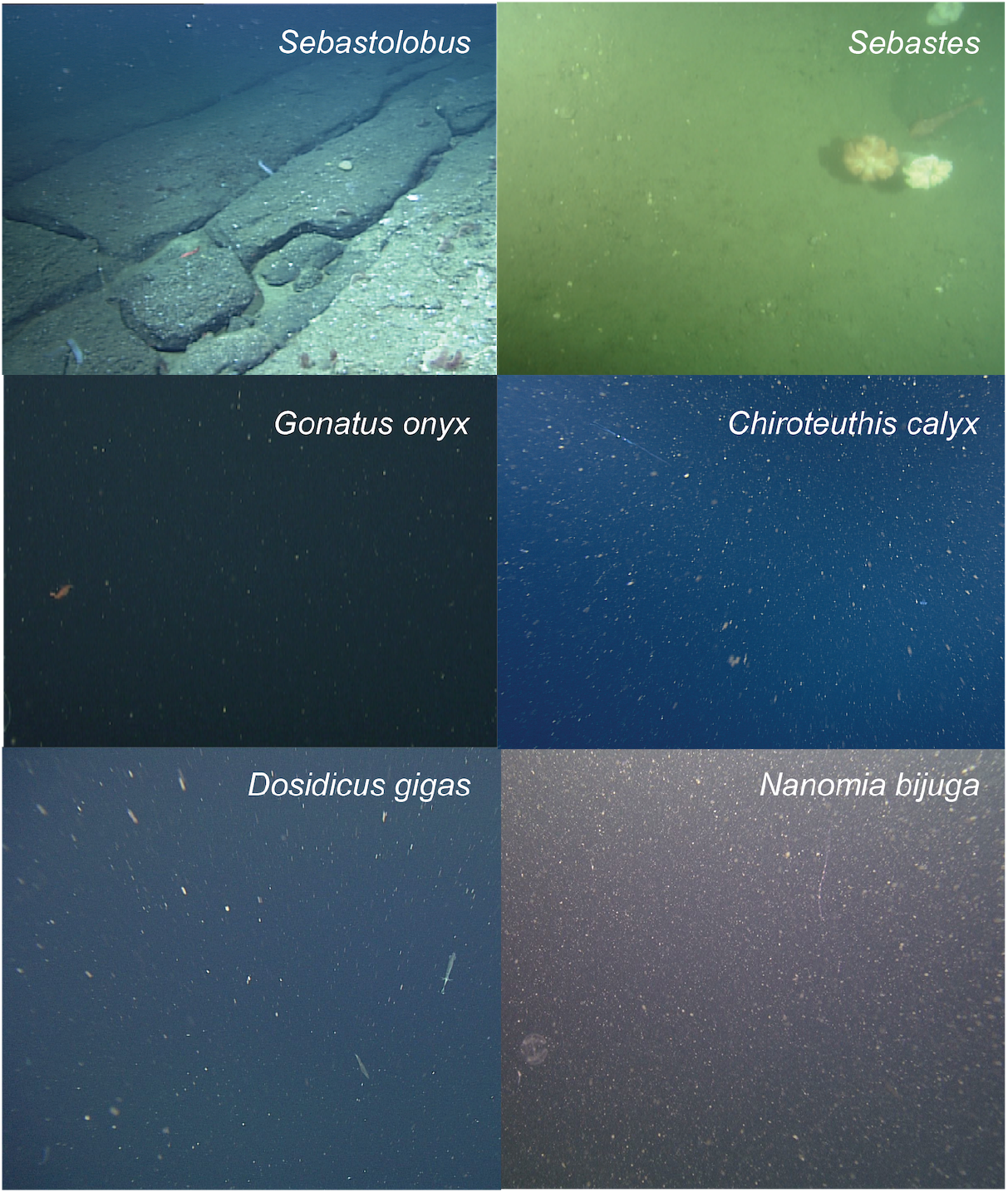
Typical images from *FathomNet* containing focal species. Focal species from Figure 1 shown in the context of more typical images from *FathomNet*. The focal class present in each image is noted in the upper right corner. Note complex and variable visual conditions, small size of objects of interest, clutter, and complex backgrounds. These conditions are typical in marine imagery collected for scientific sampling purposes.

We selected the six classes shown in Figures 1 and 2 from the much larger set of classes available in *FathomNet* based on three criteria: (i) hundreds to thousands of human-generated labels were available for each class providing us with a sufficient number of labeled instances to explore performance of ML models under different partitions of the data, (ii) images of these classes were collected over a relatively broad spatial region and/or depth range compared to many other classes in *FathomNet*, allowing us to compare performance across spatial partitions of the data, and (iii) images of these classes were collected over many years, allowing us to partition the dataset temporally to explore performance under different methodological partitions. Because searchable metadata, including depth and collection date, are included with the images in *FathomNet*, we were able to quickly create these partitions. As described in “*Diagnosing and Improving Model Performance on New Data*” below, we use these spatial and temporal partitions of the data to illustrate how ML models can fail when applied to new data, and how to diagnose and address such performance issues (Schneider *et al*. 2020). We chose to use a relatively small number of classes as opposed to using a much larger class set (e.g., as in Katija *et al*. 2022) so that we could display and compare performance measures for each class. We will return to this case study at the end of each of the following sections to provide a concrete example of each step involved in constructing and evaluating a machine learning pipeline.

## (2) Building a dataset of labeled imagery for training and evaluating models

### 2.1. Overview

The image-based ML methods that are currently most widely applied for marine science applications are based on ***supervised learning*** (Goodfellow *et al*. 2015, Cunningham *et al*. 2008). In supervised learning problems, the user provides training examples in which the desired output corresponding to a given input is specified for a set of examples. For object detection and classification problems, training data typically consist of a set of images (the image set) in which objects of interest are localized and identified by a human annotator. Labels (also sometimes referred to as “ground truths” or “annotations”) are standardized records of semantic and, in some cases, spatial information describing what is contained within the image.

To train a supervised ML pipeline to perform image analysis automatically, one needs a suitable training dataset consisting of images and corresponding labels. A researcher has two choices for acquiring the labeled data needed to train models: manually create a set of labels to be used for training, or use images and labels provided in a pre-existing database (Table 2). At present, the number of publicly available annotated datasets containing marine imagery is relatively small, and the size and spatial, temporal, and taxonomic coverage of these datasets is still rather limited. In practice, this means that researchers will typically need to create a new training dataset of annotated imagery *de novo*. This custom training set can then be used as a stand-alone training set or combined with images and labels from existing databases to fully train an ML model to carry out a specified task (Knausgård *et al*. 2021).

**Table 2.** Publicly available databases containing annotated images from marine environments. Note that some databases are actively curated and updated over time. Image and label counts are approximate and current as of October, 2022. Links are as follows: 1. https://www.kaggle.com/datasets/smaranjitghose/sea-turtle-face-detection?msclkid=2540da87b6dd11eca46690336c5e94aa 2. https://github.com/open-AIMS/ozfish 3. https://swfscdata.nmfs.noaa.gov/labeled-fishes-in-the-wild/ 4. https://fathomnet.org/ 5. https://github.com/xahidbuffon/SUIM 6. https://data.mendeley.com/datasets/n3ydw29sbz/3 7. https://www.kaggle.com/competitions/the-nature-conservancy-fisheries-monitoring/data 8. https://coralnet.ucsd.edu 9. https://www.imageclef.org/lifeclef/2015/fish 10. https://doi.org/10.13020/x0qn-y082 11. https://doi.org/10.13020/g1gx-y834 12. https://hdl.handle.net/10.1575/1912/7341, https://doi.org/10.4319/lom.2007.5.204 13. https://doi.org/10.1109/CVPR.2012.6247798 14. https://doi.org./10.17632/86y667257h.2 15. https://www.seanoe.org/data/00446/55741/ 16. https://www.kaggle.com/c/datasciencebowl 17. https://github.com/PeiqinZhuang/WildFish 18. https://www.st.nmfs.noaa.gov/aiasi/DataSets.html 19. https://www.ncbi.nlm.nih.gov/pmc/articles/PMC6176783/ 20. https://www.nature.com/articles/s41598-018-32089-8 21. https://link.springer.com/chapter/10.1007/978-3-319-24027-5_46 22. https://tinyurl.com/UNICT-Underwater-Bkg-Modeling 23. https://www.imageclef.org/lifeclef/2017/sea 24. https://www.godac.jamstec.go.jp/jedi/e/index.html

| Dataset Name | Subject | Approx. label count | Label type | Label file type | Geographic location | Published reference | URL |
| --- | --- | --- | --- | --- | --- | --- | --- |
| Save the Turtles | Turtles | 2,000 | Bounding box | .txt | Global | NA | 1 |
| OzFish | Fish | 45,000 | Bounding box | .json | Australia | doi: [10.25845/5e28f062c5097](https://doi.org/10.25845/5e28f062c5097) | 2 |
| Labeled fish in the wild | Fish | 1,000 | Bounding box | .dat | California | doi: 10.1109/WACVW.2015.11 | 3 |
| Fathomnet | Marine organisms and objects | 75,000 | Bounding box | .json | Global | doi: 10.1038/s41598-022-19939-2 | 4 |
| SUIM (Semantic Segmentation of Underwater Imagery) | Marine organisms and objects | 1,500 | Semantic segmentation | .bmp | Global | arXiv: 2004.01241 | 5 |
| Fish-Pak | Fish | 900 | Whole image | NA | Pakistan | doi: 10.17632/n3ydw29sbz.3 | 6 |
| Nature Conservancy Fisheries Monitoring | Fish aboard boats | 8,000 | Whole image | NA | Global | NA | 7 |
| CoralNet | Coral | 94,000,000 | Semantic segmentation | NA | Global | NA | 8 |
| LifeCLEF-16 Fish Dataset | Fish | 9,000 | Bounding box | .xml | Global | doi: 10.1007/978-3-319-44564-9_26 | 9 |
| Trash-ICRA19: A Bounding Box Labeled Dataset of Underwater Trash | Marine robotics, debris, fauna | 5,500 | Bounding box | .json | Sea of Japan | doi: 10.1109/ICRA.2019.8793975 | 10 |
| TrashCan 1.0: An Instance-Segmentation Labeled Dataset of Trash Observations | Marine robotics, debris, fauna | 7,000 | Instance segmentation | .json | Sea of Japan | arXiv: 2007.08097 | 11 |
| Woods Hole Plankton Dataset | Marine plankton | 3,500,000 | Whole image | NA | Woods Hole Harbor | doi: 10.4319/lom.2007.5.204 | 12 |
| Moorea labeled corals (MCL) | Corals and non-corals | 400,000 | Semantic segmentation | NA | Mo'orea | doi: 10.1109/CVPR.2012.6247798 | 13 |
| RSMAS + EILAT | Corals | 2,000 | Whole image | NA | Red Sea | doi: 10.17632/86y667257h.2 | 14 |
| ZooScan | Marine zooplankton | 19,000 | Whole image | NA | France | doi: 10.1093/plankt/fbp124 | 15 |
| Kaggle Plankton Data | Marine plankton | NA | Whole image | NA | Hatfield Marine Science Center | NA | 16 |
| Wildfish | Fish | 55,000 | Whole image | NA | Global | doi: 10.1145/3240508.3240616 | 17 |
| Labeled fishes in the wild | Fish | 1,000 | Bounding box | NA | Southern California Bight | doi: 10.1109/WACVW.2015.11 | 18 |
| DIDSON Imaging Sonar fish dataset | Fish | 1,500 | Whole image | NA | Ocqueoc River, Michigan, USA | doi: 10.1038/sdata.2018.190 | 19 |
| OBSEA EMSO | Fish, underwater scenes | 1,200 | Whole image | NA | OBSEA-EMSO testing-site | doi: 10.1038/s41598-018-32089-8 | 20 |
| FishCLEF-2015 | Fish | 14,000 | Semantic segmentation | .xml | NA | doi: 10.1007/978-3-319-24027-5_46 | 21 |
| UNICT Underwater Background Modeling dataset | Underwater scenes | 3,500 | Semantic segmentation | .xml | NA | doi: 10.1016/j.cviu.2013.12.003 | 22 |
| SeaCLEF-17 Dataset | Fish, marine animals | NA | Whole image | .xml | Taiwan | NA | 23 |
| Japan E-Library of Deep Sea Images | Organisms, geologic features, debris | NA | Whole image | NA | Deep-sea environments | NA | 24 |

### 2.2. Technical considerations

When building and working with training datasets, there are several issues a researcher should consider that can help determine which software tools are most useful, and how to best structure the labeling process to solve the desired image analysis task.

#### Label types

The most common method for creating new labels involves manual labeling of imagery (Mahajan *et al*. 2018; see Ji *et al*. 2019 for discussion of unsupervised methods). The type of label used depends on several considerations. The first consideration is the type of image analysis task that will allow the researcher to access the information they want to extract from the imagery (see “*Defining the image analysis task”* above).

If the objective is image classification, that is, to classify whole images into categories, then labels consist of a class label assigned to each image in the training set. For example, suppose the objective is to take in new images and to determine which images contain sharks (e.g., as in Sharma *et al*. 2018), and which do not. An appropriate training dataset would consist of a set of representative images from sampling cameras, each of which would be labeled by a human annotator as containing or not containing sharks.

If the objective of image analysis is to localize and classify objects within an image, then manually generated labels must contain information about the locations and classes of objects of interest within an image. The most commonly used labeling formats for object detection are bounding box labels and polygon labels. Bounding boxes are rectangular regions that enclose each object of interest and carry the appropriate class id for the object. Polygon labels, sometimes also referred to as “masks” or “segmented instances”, are enclosing polygons that outline an object of interest. These too are associated with the class label of the object. Polygon labels contain more information about the locations of the borders of each object than do bounding boxes. However, in our experience, they are significantly slower to produce, so a researcher should carefully weigh the benefits of more precisely characterizing the boundaries of objects of interest against the costs of manually labeling images with polygons rather than bounding boxes.

If the objective of image analysis is to assign the pixels in an image to distinct classes (*i*.*e*., semantic segmentation), for example to compute the fraction of the region captured in an image composed of different types of benthic cover, then labels must assign the pixels in an image to distinct classes. This is typically done within labeling software by manually selecting the borders of local regions within the image and assigning a class to these regions. Some semi-automated “assisted methods” have been developed to aid in semantic labeling of images (e.g., Uijlings *et al*. 2020).

Machine learning-based computer vision libraries such as Detectron 2 (Wu *et al*. 2019) and Deeplab v3+ (Chen *et al*. 2018) contain models that operate on an additional type of label referred to as a ***panoptic label***. Panoptic labels include both class assignments for each pixel within an image and instance labels, so that the distinct pixels belonging to an individual instance of an object, for example, an individual phytoplankton cell, are grouped together. We are not aware of past studies in marine science that have made use of panoptic labels, however, this type of labeling and segmentation could be useful in cases where a researcher wants to simultaneously characterize foreground objects of interest and background or substrate conditions.

#### Labeled data file formats

A variety of formats exist for storing manually generated labels. Unfortunately, there has been little standardization of the file formats used to encode labels of marine imagery, nor have researchers included consistent metadata within these files. When creating new labels, we recommend choosing from among several formats that are most widely used in the computer vision community, which include YOLO text files, Pascal VOC XML files, and COCO (“common objects in context”, https://cocodataset.org/) Java Script Object Notation (JSON) formats. Pascal VOC and COCO formats both allow for convenient storage of metadata, making these particularly attractive options.

#### Software for manually labeling imagery

A web search for the term “image labeling” will return many graphical user interface-based software tools designed to help users perform manual image labeling. In our experience, many of these tools work reliably, and are easy for human annotators to learn to use. Some widely-used, free labeling tools are CVAT (https://cvat.org), VGG Image Annotator (https://www.robots.ox.ac.uk/~vgg/software/via/), and Annotator J (https://biii.eu/annotatorj). Tools developed specifically for use in marine environments include BIIGLE (Langenkamper *et al*. 2017), VIAME (Richards *et al*. 2019), and EcoTaxa (Picheral *et al*. 2017; see Gomes-Pereiria *et al*. 2016 for a review). These software tools are typically intuitive to use, but different tools have different capabilities that are important to understand when deciding which package to use for a given project. When selecting a software tool, there are four issues we suggest considering: (i) the speed and ease with which images can be loaded, labeled, and the labels exported; (ii) features the labeling tool offers such as convenient batch loading of images, zooming in and out, rotating images, etc.; (iii) the label types the software allows (i.e., whole image labeling, bounding box labels, polygon labels, semantic labels, panoptic labels); and (iv) and labeled data file formats the software is capable of importing and exporting. The latter consideration is an important one, because in order to use a given set of labels to train a given ML model, the format in which those labels are stored must be compatible with the ML pipeline. For example, several freely available labeling tools only export labels in custom comma separated value (CSV) or JSON formats that are not directly readable by widely used implementations of popular machine learning architectures, including YOLO. These files can be converted from one format to another, so long as all the necessary metadata are contained within the label file, but performing such conversions introduces an additional processing step that is unnecessary if labels are exported in a suitable format.

#### Publicly available databases of annotated imagery from the field

In the computer vision literature, large, publicly available labeled image datasets such as ImageNet (14.2 million images; Russakovsky *et al*. 2015) and COCO (over 320,000 images; Lin *et al*. 2014) have been pivotal in driving the development of image-based ML methods. These datasets provide researchers with a source of data for quickly testing new model architectures, and for benchmarking and comparing new models using the same data sources. However, while large image datasets like these contain hundreds of thousands to millions of images, and a great diversity of object classes, perhaps not surprisingly, they contain relatively few images and label classes that are directly relevant to the use cases of interest to most marine scientists (Qin *et al*. 2016). Over the past decade, a number of curated open source databases containing labeled imagery from marine environments have begun to come online. The largest and most thoroughly curated of these are listed in Table 2. Depending on the specific problem a researcher is interested in addressing, these datasets may provide useful resources for model pre-training (Salman *et al*. 2016, Orenstein and Beijbom 2017, Knausgård *et al*. 2021, Li *et al*. 2022), or if classes of interest are contained within one or more of these datasets, they may contain sufficient examples to train an initial model that can be tested on new imagery and fine-tuned with new labels as necessary. As a general rule, in most marine applications, models are trained on at least some new manually labeled imagery. However, as the base of high-quality, publicly available labeled marine imagery grows, this practice may change.

#### Size of training set and balance among classes

An obvious question that arises when creating a training dataset is the question of how many images are required to achieve a desired level of performance. Several recent studies from the field have sought to address this question for the tasks of instance segmentation (Ditria *et al*. 2020) and whole image classification (Villon *et al*. 2021). In these studies, performance metrics begin to saturate between 1,000 and 2,000 labels of a given class, and adding additional labeled data from that class resulted in diminishing gains in performance. This saturation of performance around roughly 1,000 labeled instances per class is also consistent with other analyses of ML model performance on field imagery. For example, a recent analysis of the impact of label count on precision using the much larger Parks Canada terrestrial camera trap dataset reported saturation of model precision for classes with more than approximately 1,000 instances (Schneider *et al*. 2020). While the precise number of labels required to provide a desired level of performance is unlikely to follow a hard and fast rule, such numbers do provide ballpark estimates of the number of labeled instances per class one ought to have before it is worth training and testing an ML model. It is worth noting, however, that many studies that report saturating performance as label count increases compute these metrics on test sets selected at random from the overall set of images used to train, test, and validate models (e.g., Ditria *et al*. 2020, Villon *et al*. 2021, Schneider *et al*. 2020). As we will show later, the method of test set construction can have a major impact on perceived model performance.

In practice, when constructing training sets, several factors are likely to influence the number of training labels available for each class. The first is the time and cost required to manually generate labels. Whole image classification by humans can be reasonably fast (e.g., 5 seconds per image, Villon *et al*. 2018); instance segmentation tends to be slower (e.g., 13.5 sec per image, Ditria *et al*. 2020); and more elaborate label types like panoptic labeling are slower still (e.g., up to 20 minutes per image, Uijlings *et al*. 2020). How much time might it cost to create a labeled dataset? Assuming the per-image human instance labeling rate reported by Ditria *et al*. (2020), it would take 3.75 hours to label 1,000 images, which is not insignificant if objects of many different classes must be labeled. For example, to create a dataset the size of the Parks Canada camera trap dataset (47,279 images of 55 classes), a human labeler would spend roughly 21 days of full-time work (8 hours per day) labeling images. Katija *et al*. (2022) performed a more detailed valuation of the data contained within the initial release of *FathomNet* and estimated the initially uploaded dataset consisting of approximately 66,000 images to have taken over 2,000 hours of expert annotation time at a cost of roughly $165,000 for the labeling effort alone.

It clearly can be expensive to produce large labeled datasets. However, a second and perhaps a more challenging constraint has to do with limited availability of images of rare classes. Even relatively large annotated image databases from the field like the Parks Canada dataset contain many classes with far fewer than 1,000 instances per class. For example, only eight of 55 classes in the Parks Canada dataset analyzed by Schneider *et al*. (2020) are represented by more than 1,000 instances, and roughly half of all classes have fewer than 100 instances. Given the highly skewed distribution of species abundances documented in ecosystems around the world (McGill *et al*. 2007), it is simply expected that few species will be common, and most species will be far rarer. This distribution of species abundances is likely to result in image sets that contain relatively few training images of most species (Villon *et al*. 2021). Thus, we recommend constructing training datasets in a way that seeks to adequately represent all classes, but note that this may simply be impossible in many cases. In those cases, using training routines (e.g., weighted penalization of errors, Schneider *et al*. 2020; hard negative mining, Walker and Orenstein 2021) and ML pipelines that enhance performance on rare classes may be the only option. As an example of the latter, Villon *et al*. (2021) recently showed that ***few-shot learning*** models have lower asymptotic performance than more conventional machine learning models, but that the performance of few-shot learning models saturates with tens of training examples per class rather than thousands. For this reason, development of few-shot learning methods is likely to be an important area of research in the coming years.

#### Scope of training imagery versus deployment imagery

One common source of underperformance of machine learning methods on new imagery can be traced to the range of conditions and class distributions present in the training set relative to new datasets on which the model is to be used. A good rule of thumb is to try to create a training dataset that spans the range of conditions you expect to sample when deploying the model on new imagery. For example, if you are building an object detector that you intend to use in shallow water habitats and in deep water habitats, the training set should include imagery from both habitats and not just one. If you plan to use an image classifier on shallow water imagery collected from 30 distinct sampling locations across the daylight cycle, to the extent possible, train on imagery that contains the spatial and temporal variation inherent in this target use case. It is worth noting that following this advice does not necessarily mean labeling more imagery, but rather, labeling images collected from a set of conditions that better reflects the range of conditions expected when the model is applied to new image data. For instance, rather than labeling 100 images from a single sampling site at midday, one could label 10 images at each of 10 sampling sites, and choose images at each site that span the daylight cycle. González *et al*. (2017) provide a detailed discussion of strategies for building a training and validation routines that yields reliable estimates of the future performance of a trained ML pipeline.

#### Labeling errors in training data

Another factor that can degrade performance of an ML pipeline is errors in the training data. Due to human error and task ambiguity, errors in the labels used to train ML models are inevitable (Culverhouse *et al*. 2003). However, depending on how labeling was done (e.g., by a domain expert versus through a crowdsourcing platform such as Amazon Mechanical Turk; Beery *et al*. 2018), the frequency of labeling errors may vary widely (Ditria *et al*. 2020). The impact that such labeling errors ultimately have on the performance of a trained model can depend on the nature of the error. For example, in an animal pose-estimation task, Mathis *et al*. (2020) found that unlabeled objects – that is, objects of interest that were erroneously missed during the labeling process – had a lesser detrimental effect on trained model performance than did mislabeled objects – objects that were labeled, but assigned identities that were incorrect. Because correcting errors is costly, in some cases it may be worth conducting small numerical experiments that corrupt labels, for example by removing some labels or switching their class assignments. This can aid in identifying the error types of greatest concern, which can in turn inform the development of protocols for training human labelers.

### 2.3 Case study: Bounding box data from the *FathomNet* database with species- and genus-level class labels

As described above, our case study focused on six biological taxa detected in imagery collected in the Monterey Bay and surrounding regions of the coastal eastern Pacific. Images and corresponding labels for the classes used in our case study can be downloaded programmatically from *FathomNet*. Labels are downloadable from *FathomNet* in the widely-used COCO JSON format, which includes object bounding box instances corresponding to each image, along with their classes, and metadata associated with each image. Because we wished to apply a ML model called YOLO that does not accept COCO JSON as an input format, we had to convert labeled data to an admissible input format and create the necessary directory structure. Code to download images, perform conversion, and organize directories is provided at https://github.com/heinsense2/AIO_CaseStudy.

## (3) Selecting and training a machine learning model

### 3.1 Overview

After specifying an image analysis task and building a training dataset, the next step is identifying a particular machine learning model to train and test. In this review, we are focused primarily on modern computer vision methods for automated analysis, many of which rely on deep learning – learning algorithms that involve the use of ***deep neural networks*** (DNNs). However, a range of more traditional machine learning and computer vision methods have been used to perform similar tasks (Kubat *et al*. 1998, Li *et al*. 2022). Li *et al*. (2022) provide a review of traditional methods that do not rely on deep learning, and a comparison between those methods and modern deep learning methods.

### 3.2. Technical considerations

Deep learning is a form of representation learning, in which the objective is not only to use input data (e.g., an image) to make predictions (e.g., the class to which the image belongs), but also to discover efficient ways to represent the input data that make it easier to make accurate predictions (Bengio *et al*. 2013). Deep learning models are representation learning algorithms that teach themselves which features of an image are important for making predictions about the image. By training on a set of labeled images, these algorithms learn a mapping between raw pixel values and the desired output based on these features. Foundational work in deep learning demonstrated that networks that are good at representing features useful for prediction often share common structural features (LeCun *et al*. 2015), and this idea has fueled the use of deep neural networks with convolutional structure (Convolutional Neural Networks or CNNs), network pre-training (e.g., Schneider *et al*. 2020), and other practices that help ensure that networks can quickly be trained to perform a target task on a new dataset, rather than having to be fully re-designed and trained *de novo* for each new application.

#### Selecting a machine learning model

The field of DNN-based models capable of performing image classification, object detection, and semantic segmentation is enormous, and expanding by the day. Table 3 provides a list of models that have shown promising results on imagery collected from either marine environments, or terrestrial environments that present similar challenges to those frequently encountered in marine environments (e.g., complex backgrounds, heterogeneous lighting, variable image quality, *etc*.). In a practical sense, choosing which ML model to use in any particular setting involves first determining which models can perform the target task (e.g., whole image classification vs. semantic segmentation). For any given target task, there will be many available models to choose from. We recommend researchers consider three things when choosing from among these models: (i) have previous studies evaluated and compared model performance? Has any study been done that applied a particular model in a similar setting with favorable performance? (ii) Is open-source code or a GUI-based implementation of the model available? If so, how easy does it appear to be to implement? Is it compatible with the computational hardware you have available? (iii) How many additional packages, software updates, and other back-end steps are required to be able to train and deploy a given model using new data? In our experience, perhaps *the* major hurdle associated with applying any given ML model to a new dataset is the time required to configure the software and system specifications necessary to run the model code. This “implementation effort” may ultimately dictate which model an end user ultimately selects. If a given ML model has been shown to exhibit good performance, but the code available to implement that model requires significant knowledge of command-line interfaces, software package installers or dependencies, virtual environment management, hardware compatibility, or GPU programming, it may simply require too much invested time at the outset to be a viable option for most researchers. It is common practice in the deep learning and computer vision communities to provide code with published methods and applications of those methods to new datasets, and this trend is starting to be adopted by some researchers who are applying such methods to field imagery (see Table 3). However, the ease of use and level of expertise required to implement such code and interpret results varies widely from one case to another.

**Table 3.**
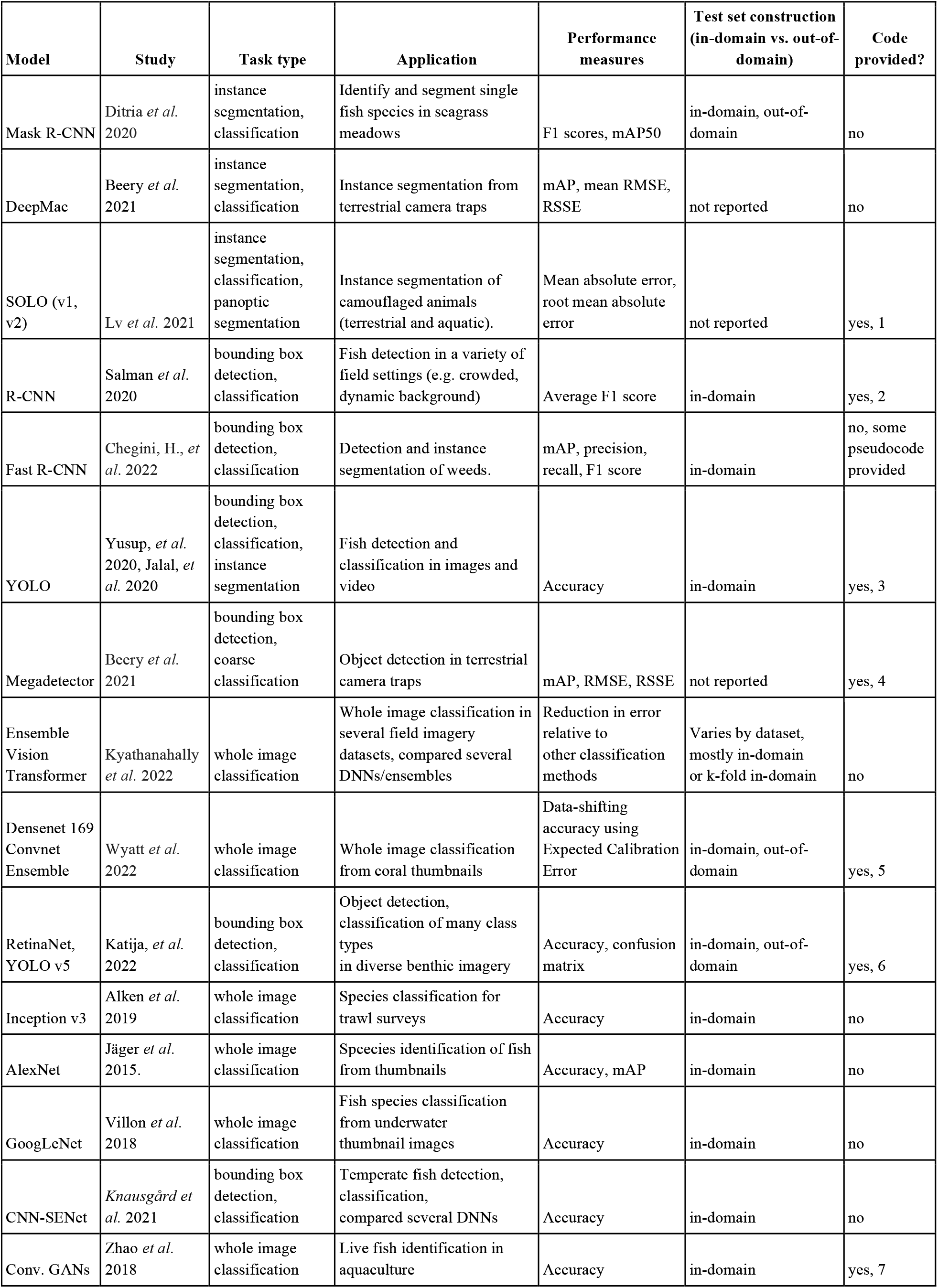
Machine learning models applied to analysis of field imagery. A selection of past models used to perform image analysis tasks on field imagery. *Performance measures reported* describes which performance measures were reported for test sets in each study. *Test set construction* describes whether the statistics reported were computed using a test set derived from the same overall dataset used to train the model (“in-domain”), or whether the test set was deliberately constructed using data from new spatial or temporal regions (“out-of-domain”). The *Code provided* column indicates whether the study provided the code used in their analyses. Code links are: 1. https://github.com/aim-uofa/AdelaiDet/ 2. https://github.com/ahsan856jalal/Fish-Abundance 3. https://github.com/ahsan856jalal/Fish-detection-and-classification-using-HOGY.git 4. https://github.com/microsoft/CameraTraps/blob/main/megadetector.md 5. https://doi.org/10.5281/zenodo.6317553 6. https://github.com/fathomnet/models 7. https://github.com/Zhaojian123/Transactions-of-the-ASABE

#### Hardware implementation: CPU vs. GPU, local vs. cloud

Another decision a user must make when implementing ML pipelines is whether to run the computations involved in training, testing, and deploying the model on a computer’s central processing unit (CPU) or on the computer’s graphics processing unit (GPU). Among the technological developments that enabled widespread use of DNN models is software and hardware innovations that allow these models to be trained rapidly and in parallel using GPUs. The technical details of ML implementations on these two distinct types of hardware are discussed in Goodfellow *et al*. (2015) and Buber and Diri (2018). The advantage of training using a CPU is that any computer can, at least in principle, be used to perform training without the need for specialized hardware that some computers have and others do not. The disadvantage is that, in the absence of custom parallelization, training a DNN model of any depth using CPUs can be prohibitively slow. Fortunately, many consumer-grade workstations now ship with GPUs that are compatible with deep learning frameworks like PyTorch (Paszke *et al*. 2019) and Tensorflow (Abadi *et al*. 2015), and many universities and research institutes are investing in shared GPU clusters. Another option for accessing machines capable of training ML models is through cloud computing services such as Google Colab, Amazon Web Services, Microsoft Azure, and others.

### 3.3. Case study: object detection and classification with YOLO

Our case study task involves detecting objects of interest, along with a bounding box and class label for each object. We selected one of the most widely used and, in our experience, robust, ML-based object detection and classification pipelines, YOLO (“You-Only-Look-Once”, Redmon *et al*. 2016). YOLO is heavily used in industry and research applications, has fast deployment times relative to other deep architectures, and is relatively easy to use. Moreover, various versions of YOLO have been incorporated into more complex detection and classification pipelines that have shown promising results on marine imagery (e.g., Knausgård *et al*. 2021, Peña *et al*. 2021). We used initial weights provided in YOLO v5 from pre-training on the COCO dataset (https://github.com/ultralytics/yolov5). We selected the “small” network size as a compromise between network flexibility and the number of network weights that need to be estimated during training. Prior to training and testing, we reduced the resolution of images to 640 × 640 px (the impact of changing resolution is evaluated below). We included the five classes of squid and fish in our primary analysis, and reserved images of the siphonophore, *N. bijuga*, for a later analysis (see “*Distractor classes*” below).

We benchmarked training and deployment of YOLO v5 using both in-house hardware (a single workstation with four GPUs), and a cloud-based implementation. For the local hardware implementation, we used a Lambda Labs Quad workstation running Ubuntu 18.04.5 LTS and equipped with four NVIDIA GeForce RTX 2080 Ti/PCIe/SSE2 GPUS, each with 11,264 MB of memory. The machine also had a 24 Intel Core i9-7920X CPUs @2.90GHz with 125.5GiB of memory. Our cloud implementation used Google Colab (https://colab.research.google.com), a cloud-based platform for organizing and executing Python programs using code notebooks (termed “Colab Notebooks”). Our cloud implementation made use of these resources using a Google Compute Engine backend with a single Tesla T4 GPU. In both local and cloud implementations, all models were trained for 300 epochs (or for fewer epochs when early stopping conditions were met) using all available GPUs. Run times on our local and cloud implementations were comparable, with the 4 GPU local machine performing slightly faster (mean of 19.1 sec per training epoch; 1.59 hours to complete 300 epochs) than the single GPU cloud implementation (mean of 26.8 sec per training epoch; 2.23 hours to complete 300 epochs).

## (4) Evaluating model performance

### 4.1. Overview

After training models, a final step in the model building process is to evaluate model performance. Many metrics are available for measuring the performance of ML models, and the most appropriate metric in any given application will depend both on the task the model is trained to execute (e.g., image classification vs. semantic segmentation), and the relative importance of different kinds of errors the model can make (e.g., false positives vs. false negatives), which must, of course, be determined by the researcher. Goodwin *et al*. (2022) and Li *et al*. (2022) provide approachable discussions of common metrics, along with formulae for computing them and the logic that underlies them. Tharwat (2020) provides a more technical account of classification metrics and their strengths and weaknesses. In very general terms, one typically wishes to evaluate the ability of the ML model to predict the correct class of an object, image, or subregion of the image, and, if the method provides spatial predictions about objects or semantic classes located in different parts of the image, one would like to know how accurate these spatial predictions are.

### 4.2. Technical considerations

For whole image classification, performance metrics seek to express the tendency of the model to make different kinds of errors when predicting classes. For example, suppose a researcher has 300 sea surface satellite images, and a model is trained to determine which images contain harmful algal blooms (HABs) and which do not (Henrichs *et al*. 2021). The ***classification accuracy*** of the model is the ratio of images that were assigned the correct class (HAB present vs. HAB absent) over the total number of images classified: (true positives + true negatives) / (total images classified). If the model correctly predicted 100 images that contained HABs, and correctly predicted 100 images that did not contain HABs, the accuracy is 200/300 = 0.67. Accuracy is an appealing measure because of its simplicity but there are many cases in which it can be misleading, particularly when the dataset contains multiple classes and the relative frequency of classes differs (see discussion in Tharwat 2020). Other widely-used metrics including precision, recall, and F1 score, were designed to capture other aspects of model performance, while avoiding some of the biases of classification accuracy. The ***precision*** of a classifier measures the fraction of positive class predictions that are correct. If the model classifies 130 images as containing HABs and 100 of these images actually contain HABs, the precision of the classifier is 100/130 = 0.77. ***Recall***, sometimes also referred to as “sensitivity,” measures the ability of a model to detect all images or instances of a given class that are present in the dataset, thereby expressing how sensitive the model is to the presence of a class. If the classifier correctly classifies 100 images containing HABs but the dataset contains 160 images that contain HABs, the model’s recall is 100/160 = 0.63. The ***F1 score*** provides a composite performance measure that incorporates both precision and recall: *F1 = 2 (precision x recall)/(precision + recall)*.

For methods that make spatial predictions, such as object detection or semantic segmentation, there is an additional question of whether the model’s spatial predictions are located in the right place. Among the most widely-used methods for measuring the spatial overlap between predictions and data this involves computing the spatial overlap between a prediction from the model and objects in the labeled image. This is often measured using the ***intersection-over-union*** (IoU): the intersection area of the predicted borders or bounding box of an object and the borders or bounding box of the label, divided by the total number of unique pixels covered by the bounding box and the label. Pairs in which the predictions precisely overlap labels will have equal intersection and union, giving an IoU value of one. Complete mismatches, partial spatial matches, or cases where the predicted and labeled bounding boxes differ in size will result in a union that exceeds the intersection and an IoU value less than one, with a minimum of zero in cases where there is no overlap between predicted and observed bounding boxes.

In object detection and classification tasks, the added complication of predictions being spatial raises some questions about how one ought to compute the accuracy of class predictions. A standard practice is to consider a given bounding box a valid “prediction” if its IoU value exceeds some pre-specified threshold, which is often set arbitrarily at 0.5. For bounding box-ground truth pairs exceeding this threshold, one then evaluates performance using the same types of methods applied in whole image classification (e.g., accuracy, precision, recall, f1 score, etc.). A widely-used measure is known as the ***mean average precision*** (mAP), which is most commonly calculated from the precision-recall curve as the average precision of model predictions over a set of evenly spaced recall values between zero and one (Everingham *et al*. 2010), where the precision-recall curve represents model precision as a function of model recall across a range of values of a threshold parameter. The thresholds most often used are the box or instance confidence score and the IoU of predicted and labeled object detections. By default, YOLO v5 produces two measures of mean average precision: mAP@0.5, which is the mean average precision of the model assuming matches constitute all prediction-ground truth pairs with IoU >= 0.5, and a second measure, mAP@0.5:0.95, which is the arithmetic mean of average precision of the model computed across a range of threshold IoU values in the set {0.50, 0.55,0.60,…,0.90, 0.95}. It is worth noting that different studies and machine learning software implementations compute mAP slightly differently, so ensuring that you understand how it is being computed in any given instance is important when comparing predictions across studies or ML methods.

#### Cross validation and performance evaluation

When evaluating the performance of a model on test images that were held out during training, the exact values of performance metrics will depend on the particular subset of images used during testing. Because training, validation, and testing image sets are typically selected at random from the overall image set (e.g., see Table 3), in principle, they should reflect the overall conditions and variability in the parent dataset. However, random variability in exactly which images end up in training, validation, and test sets will invariably introduce stochasticity in performance estimates. One way to address this is to create several or even many random subsets of the overall image dataset into training, validation, and test sets (Schneider *et al*. 2020). This is sometimes referred to as ***k-fold cross validation***, where *k* denotes the number of training/validation/test splits included in the analysis. The objective of this type of cross validation is to provide more robust measures of performance by averaging over multiple random partitions of the data into training, validation, and testing sets.

#### Non-random partitioning and “out-of-domain” performance

In addition to cross validation using random partitions of the data, it is also becoming more common to evaluate model performance on non-random partitions of data into training/validation and test datasets (Taori *et al*. 2020, Schneider *et al*. 2020). Typically, this is done to produce test sets that are more representative of new data on which the ML pipeline is intended to be used. For example, if one wishes to train an image classifier to classify coral species from images (Wyatt *et al*. 2022), and this classifier is intended to be used at new locations in the future, one way to test its performance would be to divide the annotated imagery available into distinct spatial locations, and to construct the training and validation set from a subset of those locations, while holding out other locations that the model never sees during training. This type model evaluation seeks to determine whether models are capable of performing well on images that may have very different statistics than the images on which they were trained. We will come back to this issue in the following section.

### 4.3. Case study: performance on object detection and classification of underwater imagery

#### In-domain performance on test imagery

Images of our target classes in *FathomNet* were collected at many different physical locations, and over decades of sampling (32 years spanning 1989-2021) using remotely operated vehicles equipped with a range of different types of imaging equipment. This led to an image set with complex and diverse backgrounds, highly variable visual conditions, and a wide range of image statistics (Fig. 2). Nevertheless, after training YOLO v5, we were able to achieve high object detection and classification performance on test imagery selected at random from the same spatial region or temporal period used to build the training set (Table 4 “in-domain”). Mean average precision (mAP) of model predictions ranged from 0.67-0.95, and three classes had mAP values of 0.88 or above. Model F1 scores had an average value of 0.77, and three classes had F1 scores of 0.81-0.92. To put these performance metrics in context, Ditria *et al*. (2020) quantified the ability of citizen scientists and human experts to detect and classify a single fish species of interest (*Girella tricuspidata*) in images taken from shallow-water seagrass beds in Queensland, Australia. Citizen scientists and experts had mean F1 scores of 0.82 and 0.88, respectively. Comparing performance of YOLO v5 on our dataset to these benchmarks implies that our detection and classification results are in the same range as those of human annotators on a similar task.

**Table 4.**
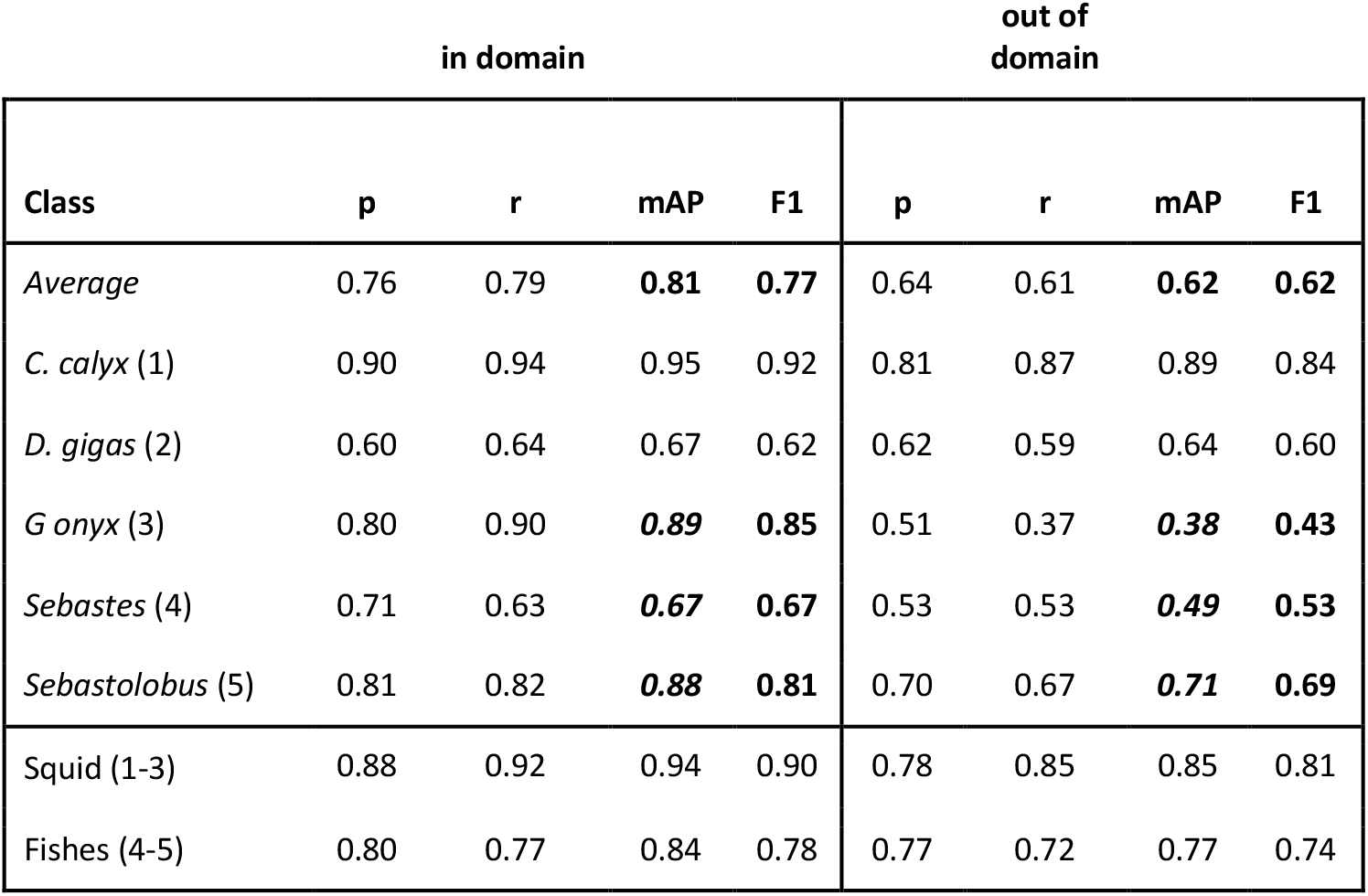
Average performance of YOLO v5 object detection and classification on images selected at random from the same spatial or temporal partition used to build the training set (“in-domain”), or the partition held out (“out-of-domain”). Note near universal decrease in all performance measures in out-of-domain data consistent with distribution shifts across spatial and temporal partitions. Drops in mAP and F1 between in-domain and out-of-domain sets of greater than 0.10 are bolded. “Squid” and “Fish” rows give results for class coarsening experiment (see “*Class Coarsening*” in text), where species and genus-level classes are aggregated into coarser classes, fishes (*Sebastes* and *Sebastolobus*) and squid (*C. calyx, D. gigas*, and *G. onyx*). Note mAP and F1 scores on “Fishes” class in out-of-domain data exceeds performance on either of the individual fish genera, indicating an overall enhancement in performance through class aggregation.

#### Out-of-domain performance: evidence for distribution shifts

As noted above, many researchers who wish to use machine learning pipelines to analyze imagery from the field often intend to use trained pipelines to analyze *new* imagery taken at later dates or different physical locations, rather than focusing solely on images taken from the same database used to construct the training set (Wyatt *et al*. 2022, Beery *et al*. 2018, Schneider *et al*. 2020). To simulate this scenario, we performed a nonrandom, four-fold cross-validation procedure on the overall set of annotated imagery available on our classes of interest in *FathomNet*. This involved two different kinds of nonrandom partitioning of the dataset. The first was a temporal partition, in which we divided all annotated images of our focal classes into images collected prior to 2012, and images collected from 2012 through the present. This partitioning resulted in *pre-2012* and *post-2012* (2012 onward) image subsets. Splitting the data at 2012 yielded a similar number of labeled instances for most classes before and after the split. The second partition we performed was a spatial partition. Images from all sampling dates were pooled together. But for each class, we divided images either by depth or by latitude and longitude to ensure that images of each class were divided into distinct spatial “regions,” defined arbitrarily as *region 1*, and *region 2*. This temporal and spatial partitioning resulted in a four-fold partition of the data: two temporal sampling periods, and two spatial regions. We measured average performance over the four data partitions by training on one of the partitions and testing on the other (e.g., training on *pre-2012* images and testing on *post-2012* images, training on *region 1* and testing on *region 2*).

Figure 3 and Table 4 shows the results of this analysis. Performance metrics were generally lower in the out-of-domain partition than in the partition from which training data were drawn (general trend of decreasing performance evident in Fig. 3). This decrease in performance was particularly extreme for certain classes. For example, average mAP and F1 scores for the black-eyed squid, *Gonatus onyx*, were cut approximately in half – from 0.89 and 0.85, respectively, to 0.38 and 0.43 – when a model trained on one partition was deployed on the other. As previously suggested (Katija *et al*. 2022), these findings imply that ***distribution shift*** occur in the *FathomNet* dataset, and that these shifts can significantly degrade performance when models are trained on data from one set of locations or time periods and deployed on imagery from new locations or time periods. This phenomenon appears to be widespread in imagery collected in the field (Schneider *et al*. 2020, Wyatt *et al*. 2022).

**Figure 3.**
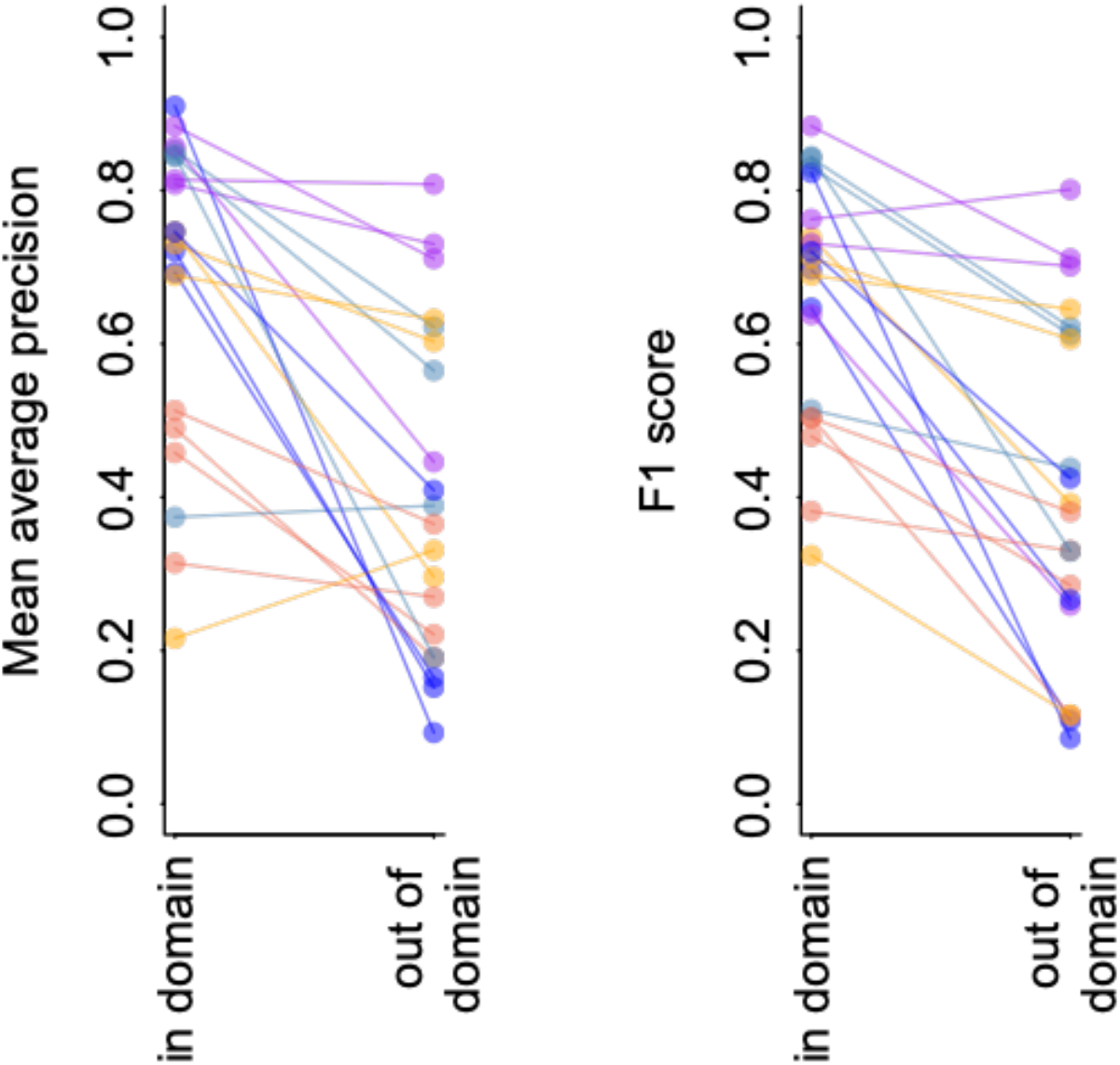
YOLO v5 model performance on imagery from *FathomNet*. Change in mean average precision (left) and F1 score (right) when a model is tested using out of sample data from the same spatial or temporal partition from which training data was selected (“in-domain”), and when the same model is tested using data from a different spatial or temporal partition (“out-of-domain”). Colors indicate different classes, and lines connect points from the same spatial or temporal partitioning of the data to indicate trends.

## (5) Diagnosing and improving model performance on new imagery

### 5.1. Overview

Although ML-based frameworks have shown impressive classification performance on imagery from marine systems (e.g., ensemble vision transformers with accuracies in the range of 96.5-99.9% for image classification tasks, Kyathanahally *et al*. 2022), inevitably, all models make errors. Moreover, the degree to which a previously trained model makes errors when applied to new image datasets can change over time as new imagery changes relative to the original dataset used to perform training. Therefore, one key step in building and maintaining a ML pipeline for automated image analysis is diagnosing performance problems and finding ways to fix them (Norouzzadeh *et al*. 2018; Schneider *et al*. 2020). In this section, we address issues that can degrade performance of an ML pipeline, and suggest approaches for remedying these issues. Many such issues can be traced back to the problem of distribution shift (Taori *et al*. 2020, Beery *et al*. 2018, Schneider *et al*. 2020). A distribution shift occurs when the imagery on which an ML pipeline is trained differs in some systematic way from the imagery on which the pipeline is deployed – that is, the new imagery the ML method is being used to analyze. The term “distribution shift” refers to a generic set of differences that may occur between one set of images (the “in-domain” set) and another (the “out-of-domain” set), including things like differences in lighting, camera attributes, image scene statistics, background clutter, turbidity, and the relative abundances and appearances of different classes of objects (Taori *et al*. 2020, Scoulding *et al*. 2022, Wyatt *et al*. 2022). This problem arises often in scientific sampling, because in this setting, the goal is often to extract data from one image set collected at a particular place and time, and to compare it to data from another image data set collected at some other place or time. Because conditions and the class composition of objects of interest can vary widely in space and time, it is likely that an image set collected at a given set of locations over a given time period will differ systematically from a new set collected at different locations or different times (Schneider *et al*. 2020). To allow for meaningful comparisons across these conditions, it is crucial that an ML model has similar performance across conditions or that any biases in performance be known when interpreting model results. If this is not the case, the ML model itself can introduce bias into measurements that can confound scientific comparisons between image datasets collected from different locations or time periods.

Many existing ML methods for object detection and classification perform poorly under distribution shifts without careful training interventions (Beery *et al*. 2018, Taori *et al*. 2020, Schneider *et al*. 2020). Despite this, human labelers exhibit similar performance on original and distribution shifted datasets (Shankar *et al*. 2020), suggesting that distribution shifts do not reduce the information needed to accurately identify objects *per se*, but rather that the structure and training of ML models cause them to fail on distribution shifted imagery (Taori *et al*. 2020). Given that distribution shifts are documented here (Fig. 3, Table 4), and in past studies of imagery from the field (e.g., Beery *et al*. 2018, Schneider *et al*. 2020, Ditria *et al*. 2020, Katija *et al*. 2022), a natural question is whether there are steps that can be taken to reduce the effects of distribution shifts on model performance.

### 5.2. Technical considerations

A wide array of methods have been proposed to improve the performance of ML models on new imagery that is distribution shifted relative to training images. These range from training interventions like digitally altering (*i*.*e*., *“augmenting”*) training imagery to destroy irrelevant features that can result in overtraining (Busalaev *et al*. 2020, Zoph *et al*. 2020, Bloice *et al*. 201), to the use of more robust inference frameworks such as ensemble models, which combine predictions of multiple machine learning models (Wyatt *et al*. 2022). To provide a sense for how some of these methods work, and how one might apply them to improve model performance, we applied a suite of training interventions to our case study dataset.

***Image augmentation*** is a widely used method for improving model performance on out-of-sample and out-of-domain imagery (Busalaev *et al*. 2020, Zoph *et al*. 2020, Bloice *et al*. 2019). Image augmentation involves applying random digital alterations of training imagery during the training process to help avoid over-fitting ML models to specific nuances of training imagery that are not useful for identifying objects of interest in general. Augmentation of training images is used by default in many ML pipelines (including YOLO v5) as part of the training process, but augmentation parameters are often tunable, so having some understanding of how different types of augmentation affect performance on field imagery is useful.

Increasing ***image resolution*** is another straightforward training intervention. Due to the computational and memory demands of training DNN-based ML models, it is common to reduce image resolution during training, testing, and deployment (e.g., Kyathanahally *et al*. 2022, Schneider *et al*. 2020). However, if objects of interest constitute relatively small regions of the overall image (e.g., Fig. 2), reducing resolution can coarsen or destroy object features that can be important for detection and classification. The loss or degradation of these features during training and deployment, mean that they cannot be used to accurately detect and classify objects in new imagery that is distribution shifted relative to the training set. It may, therefore, be beneficial in some applications to maintain higher image resolutions during training and deployment.

Training using ***background imagery*** is another straightforward training intervention that is relatively easy to implement. While it can be costly to label new imagery for the reasons discussed above, it can be relatively cheap to identify “background images,” defined simply as images that do not contain objects of interest. Training a ML model including background imagery in the training set has been proposed as one method for helping models to better generalize to new image sets (Villon *et al*. 2018).

A fourth type of intervention is known as ***class coarsening***. Intuitively, objects that are visually similar are likely to be harder to discriminate than are objects that look very different. Given this, one potential solution to improve model predictions under distribution shifts is to coarsen class labels in a way that results in similar looking classes being aggregated into a single super-class (Williams *et al*. 2019, Katija *et al*. 2022). In biological applications, this may result in aggregating classes with finer phylogenetic resolution (e.g., species or genus-level classes) into classes with coarser resolution (e.g., family or order-level classes or coarse species groups). For instance, rather than requesting individual species of sea fan and corals, one might simply specify “sea fans” and “corals” as classes. Whether this is a suitable training intervention will obviously depend on the ultimate goal of the image analysis and whether coarser class labels are acceptable.

A final intervention we consider is training on images that include objects in distractor classes. The definition of the term “distractor” in the computer vision literature has varied (e.g., see Das *et al*. 2021 vs. Zhu *et al*. 2018). Here, we define a distractor class as a class of object that shares visual characteristics with a target class and could reasonably be confused with the target class during classification. This working definition is consistent with the way the term “distractor” is used in the visual neuroscience literature (e.g., Bichot *et al*. 1999). When training ML models to detect a certain class or small set of classes, it is common to train models using labels of only the class or classes of interest. However, if distractor classes are regularly present in new imagery, they can degrade model performance. Deliberately including images of distractor classes in the training set is a form of adversarial training that may improve model performance when distractor classes occur in new imagery.

In addition to these relatively simple training interventions, a variety of other solutions to improve model robustness on new imagery have been proposed. These include the use of ensemble models, where predictions are derived not from just one deep neural network, but from many networks whose predictions are combined to make an overall class prediction (Wyatt *et al*. 2022), adversarial training, sometimes also called “active learning,” in which models are re-trained with images on which they previously made errors (Mathis *et al*. 2020), training on synthetic data (Schneider *et al*. 2020), and stratified training in which the relative abundance of classes in the training set are modified by excluding or including extra examples of one class or another (Schneider *et al*. 2020). There are related methods that seek instead to analyze the output of automated systems at the sample level, rather than the individual level, to correct errors and detect changes in new domains (González *et al*., 2019, Walker and Orenstein, 2021). We refer the reader to the research cited in this section, and to Taori *et al*. (2020), Santurkar *et al*. (2020) and Koh *et al*. (2021) for further reading on methods for improving performance under distribution shifts.

### 5.3. Case study: training interventions and performance on out-of-domain imagery

#### Image augmentation

To test whether and how augmentations might improve model performance on new imagery, we applied three kinds of augmentation to images during training: orientation augmentations, in which the training image and corresponding bounding box is scaled or flipped by a random amount, color space augmentations, in which the color attributes of the training image are randomly perturbed during training, and mosaic augmentation, in which sets of training images from the training set are randomly selected, cropped, and recombined to form a new composite “mosaic” image used in training. We tested the impact of each of these augmentation types by starting with all of them active, then dropping one augmentation type at a time. For each of these augmentation “treatments,” we computed the average decrease in mean average precision from the in-domain testing set to the out-of-domain testing set, averaging over all four partitions of the data.

Applying no augmentations at all resulted in the largest drop in performance from in-domain to out-of-domain imagery (an average drop in mAP of 0.27 across classes). The smallest mean drop in performance over all classes occurred when all augmentations were applied (an average drop in mAP of 0.18 across classes). However, the effects of augmentations were highly variable among different partitions of the data, and among classes. For example, the improvement in out-of-domain *mAP* from no augmentation to the most effective augmentation scheme varied by a factor of three, from 0.08 for rockfish (*Sebastes*) to 0.24 for the black-eyed squid, *Dosidicus gigas*. These results suggest that augmentation may indeed be a way to improve generalization on new imagery in datasets similar to that used in our case study, but that effects of augmentation may be highly variable from one class to another.

#### Image resolution

To explore the impact of changing image resolution in our case study, we modified the default resolution specified in YOLO v5 (640 px x 640 px) to a higher resolution (1280 px x 1280 px). Between 91% and 98% of images available in *FathomNet* for each class have a resolution equal to or greater than 640 pixels along at least one axis. Images with resolution lower than 1280 × 1280 were loaded at full resolution and padded at the borders to reach the desired training resolution. Effects of increased image resolution were not large. For example, the average change in mean average precision on out-of-domain data across the four partitions was 0.04, and the largest performance increase was only 0.05 (for *G. onyx*), while performance on *C. calyx* actually dropped slightly when we used higher resolution imagery. It is worth noting that differences in resolution between training and testing data can cause degraded performance, which may have contributed to a lack improvement in performance in some of the partitions (e.g., *pre-2012* vs. *post-2012* splits, for which image resolution systematically differed).

#### Training on background imagery

To test whether training on background imagery could improve out-of-domain performance, we re-trained YOLO v5 using the *post-2012* partition as a training set, but we also included background images from the *pre-2012* and *post-2012* partitions in the training imagery. Including background imagery improved performance on all classes (Table 5), with marked increases in performance for *Dosidicus gigas* and *Gonatus onyx*, the classes with the fewest labels in the training set (*n* = 42, and *n =* 84 labeled instances, respectively).

**Table 5.** Effect of including background imagery on performance of YOLO v5. “No BG images” shows performance of standard training in which no background images are included in the training set. “BG images” shows statistics for training runs in which background images from pre-2012 and post-2012 periods that did not contain any classes of interest were included in the training set. Bolded mAP and f1 score values in “BG images” show cases where these statistics improved relative to training without background images. Results from the two temporal partitions are averaged.

|  |  | No BG images |  |  |  | BG images |  |  |  |
| --- | --- | --- | --- | --- | --- | --- | --- | --- | --- |
| class | num. labels<br>in training set | p | r | mAP | f1 | p | r | mAP | f1 |
| *C. calyx* | 351 | 0.84 | 0.88 | 0.90 | 0.86 | 0.87 | 0.88 | 0.90 | **0.88** |
| *D. Gigas* | 229 | 0.48 | 0.46 | 0.51 | 0.47 | 0.57 | 0.60 | **0.59** | **0.58** |
| *G. onyx* | 94 | 0.66 | 0.53 | 0.57 | 0.59 | 0.76 | 0.51 | **0.60** | **0.61** |
| *Sebastes* | 445 | 0.67 | 0.54 | 0.61 | 0.60 | 0.65 | 0.60 | **0.63** | **0.62** |
| *Sebastolobus* | 1178 | 0.74 | 0.82 | 0.83 | 0.78 | 0.76 | 0.82 | **0.84** | **0.79** |

#### Class coarsening

To explore whether class coarsening improved performance under distribution shifts, we coarsened class labels from the species (*Gonatus onyx, Chiroteuthis calyx*, and *Dosidicus gigas*) and genus level (*Sebastes* and *Sebastolobus*) to the coarse categories of *squids* and *fishes*. Table 4 shows performance of YOLO v5 when trained and tested on these coarser classes. As expected, coarsening classes resulted in a smaller average drop in model performance when models were applied to out-of-domain data. For the *squid* class, out-of-domain performance was higher than for any individual class in the fine class model except for *C. calyx* (class for which the model had the highest performance). Out-of-domain performance for the *fish* class was higher than performance on either of the individual fish genera in the analysis where genera were treated as separate classes.

#### Training with distractor classes

To quantify the impact of training with distractor classes on model performance, we restricted our analysis to two classes: the swordtail squid, *Chiroteuthis calyx*, and the siphonophore, *Nanomia bijuga*. In particular, we sought to determine whether a trained ML model could discriminate images of juvenile swordtail squid in an image set containing images of juvenile *C. calyx* and imagery of *N. bijuga*, a distractor class that is a mimicked both morphologically and behaviorally by juvenile *C. calyx* (Burford *et al*. 2015). Because the spatial distributions and habitat use of these two species overlap, a researcher interested in *C. calyx* would likely need to contend with images containing *N. bijuga*, either by itself or in the same image as the target class *C. calyx* (e.g. as in Fig. 1F). A naïve approach for training a ML model to detect juvenile *C. calyx*, would be to train only on images of this target class, and then to deploy the model on new images containing one or both of the two classes.

Table 6 shows performance of YOLO v5 trained to detect juvenile *C. calyx* using this naïve approach. Mean average precision is relatively poor as are precision and recall scores (e.g., mAP = 0.54). Moreover, 22% of the instances of *N. bijuga* in test data were erroneously classified as *C. calyx*, indicating that the model often mistook the distractor class for the target class. To determine whether training on both the target and distractor class could help remedy this issue, we re-trained YOLO v5 with a training set containing labeled imagery of both juvenile *C. calyx* and *N. bijuga*. We then applied this model to test imagery. Training on both classes resulted in a pronounced increase in all performance metrics to levels that match or exceed reported performance of human labelers in similar tasks (precision = 0.86, recall = 0.9, mAP = 0.87). Moreover, despite the strong morphological resemblance between *N. bijuga* and juvenile *C. calyx* (Fig. 1F), the model trained on both classes never classified new images of *C. calyx as N. bijuga* or vice versa (0% misclassification rate). A third approach to improving model performance in the presence of distractor classes that is less costly than manually labeling distractor classes is to include in the training set images that contain the distractor, but to treat these as unlabeled “background” imagery. That is, if an image contains only the distractor class, it would be included in the training set with no instance labels. The logic behind this is to ensure that the model is trained on many example images that indicate the distractor is not the target class. Training in this way saves time because it does not require manual labeling of the distractor class or classes. To test this approach, we used the same images of *C. calyx* and *N. bijuga* used to train the two-class model, but we included no labels for the *N. bijuga* class. Performance using this approach was only slightly lower than performance of the model trained on labels of the distractor (Table 6), indicating that such training be a viable alternative to building a full dataset containing labels for distractors as well as the target class.

**Table 6.** Effects of distractor class on model performance. Detection of an object class of interest – in this case, juveniles of the swordtail squid, *C. calyx* – in imagery containing the class of interest and a distractor class, *N. bijuga*, that closely resembles the target class (see Fig. 1F). Precision (p), recall (r), mean average precision (mAP), and F1 score are shown along with misclassification frequency, the fraction of labeled instances of the distractor class, *N. bijuga*, that were erroneously classified as the target class, *C. calyx*.

|  | **p** | **r** | **mAP** | **F1** | **Misclassification frequency** |
| --- | --- | --- | --- | --- | --- |
| Train without distractor | 0.49 | 0.68 | 0.54 | 0.56 | 0.22 |
| Train with distractor as background | 0.88 | 0.70 | 0.79 | 0.78 | 0.008 |
| Train with distractor labels | 0.86 | 0.90 | 0.87 | 0.89 | 0 |

#### Summary of training interventions and their effects on performance on new imagery

Overall, we found that training on background imagery, and class coarsening provided improvements in performance on new imagery in all or most classes in the dataset. Training distractors also resulted in large improvements in performance. The impact of image augmentation was more variable, but still resulted in overall improvements in performance for all classes. Training and deploying the model on high-resolution imagery (as opposed to images with reduced resolution) had the most variable effect on performance, but this should be taken with the caveat that our image set consisted of a mix of high- and low-resolution imagery, and that resolution mismatches between training and testing data can sometimes result in poor performance.

The image set and number of classes used in our case study was intentionally limited, so our findings should also be taken with this in mind. Nevertheless, our analysis suggests that training on background imagery, distractor classes, and augmentations may be effective ways to improve performance on new imagery without changing the structure of classes. If the nature of the problem doesn’t require that classes be discriminated with high resolution, coarsening the definition of classes could also be a highly effective way to improve the ability of models to generalize to new image sets (Katija *et al*. 2022).

## Conclusions and future prospects

Image-based machine learning methods hold tremendous promise for marine science, and for the study of natural systems more generally. These methods have the potential to vastly accelerate image processing, while also greatly lowering its costs (Norouzzadeh *et al*. 2018, Katija *et al*. 2022). In doing so, these methods could fundamentally change the spatial coverage and frequency of sampling achieved by research and monitoring efforts. Our objective in this work has been to provide a guide for researchers who may be new to these methods, but wish to apply them to their own data. If image-based machine learning methods are to be more widely adopted and fully exploited by marine scientists, the current high barrier to entry associated with these methods must be lowered. We therefore conclude with four suggestions for the research community that we believe could go a long way toward expanding use of, and access to image-based machine learning tools across marine science. These suggestions are aimed at both marine scientists who collect imagery from the field, and at the community of developers currently building ML methods and the software tools necessary to implement them.

### (1) Open sharing of labeled image datasets from the field

At present, the ability of researchers to test and engineer ML methods relevant to the tasks marine scientists want to perform on imagery is hampered by a relative shortage of publicly available data for training and testing these methods. Thus, among the most important steps that can be taken to improve ML models for use in the marine domain, is to increase the availability, coverage, quality, and size of domain-relevant labeled image datasets. As Table 2 shows, available datasets focus rather heavily on tropical fishes, coral, and marine phytoplankton, whereas imagery of other kinds of objects of interest and imagery from other habitats is not as well represented. Research groups around the world are increasingly relying on imagery as a primary data source for capturing information about the ocean. Yet, most of this imagery is still analyzed manually by human experts. Researchers who generate such datasets in the course of their work would contribute much to the community by making those datasets available in a form that is easily readable by ML pipelines. This can be done either through stand-alone publications of such datasets (e.g., Saleh *et al*. 2020, Ditria *et al*. 2021), publication of datasets as part of standard research publications (e.g., Sosik & Olson 2007), or by contributing datasets to open image repositories such as *FathomNet* (Katija *et al*. 2022) and *CoralNet* (Williams *et al*. 2019). Of course, constructing labeled image datasets requires funding, domain expertise, and a significant commitment of personnel time. It is therefore crucial that researchers who generate such datasets and the funding sources that support them receive credit. Part of this process will involve a shift in perspective from viewing annotated imagery as simply a means to an end, to viewing these kinds of datasets as valid research products in their own right (Qin *et al*. 2016, Koh *et al*. 2021, Ditria *et al*. 2021). Fortunately, this shift in perspective is already beginning to occur, and we expect funding agencies, tenure and promotion committees, and the community of marine scientists more generally will continue to move in the direction of recognizing the value of producing high-quality labeled image datasets and making those datasets public.

### (2) Sharing of open source code for repeating analyses

A second recommendation is aimed at researchers who are developing and testing ML methods for analyzing imagery from the field. It is now commonplace among the larger computer vision community for preprints, conference publications, and journal publications to include links to code repositories that contain the code necessary to repeat the analyses described in the paper. We encourage researchers who are developing ML methods to solve problems in marine science to follow this same practice. Providing the code that accompanies work described in publications can accelerate research. Doing so allows others to work through the methods using the paper and code as complementary descriptions of analyses. Providing code also helps catalyze a broader application of the model across the research community, and provides a starting point for researchers to extend and improve methods. While newer studies are beginning to follow this practice, it is still not as widespread among researchers working in marine science as it is in the broader computer vision community (Table 3). Code can be efficiently shared, for example, through GitHub repositories or through “model zoo” features of existing image repositories such as *FathomNet* (https://github.com/fathomnet/models).

### (3) Develop and adopt standards for model evaluation that accurately capture performance in common use-cases

At present, there has been little standardization of model performance metrics reported in papers that apply image-based machine learning to problems in marine science. Different papers report different metrics that often include just one or a few of the performance measures described in “*Evaluating model performance*” above. It is hard to make comparisons between methods when different research groups evaluate alternative methods using different performance metrics. The most commonly reported metric across studies is classification accuracy (Table 3), but, as noted above, this metric is subject to biases that inherently make comparisons across studies problematic (Tharwat 2020). Another less obvious issue is that different studies compute standard metrics such as accuracy, precision, recall, and F1 score in different ways. For example, some studies compute performance measures from a single random partition of the data into training, testing, and validation sets. Others perform several random partitions of the overall dataset using a k-fold cross-validation procedure. Others still report true out-of-domain statistics computed on test data from specific locations or time periods that were held out during training (see Table 3). As we illustrate in our case study, the way in which test imagery is selected (e.g., at random from in-domain data vs. from out-of-domain data) can have a major impact on performance measures, and any fair comparison between methods clearly requires that performance statistics of competing methods be computed in as similar a manner as possible.

In the end, the most appropriate performance measures will be the ones that best reflect how a model will perform at the task for which it is ultimately intended to be used (González *et al*. 2017). At the same time, adopting a standard will likely be necessary if performance is to be compared among studies. To achieve this compromise, we suggest that it will be productive for the community of developers and users of image-based machine learning methods in the marine domain to begin a conversation about the most appropriate standards for evaluating models and comparing model performance among studies, with the goal of identifying metrics that meet the needs of researchers.

### (4) Develop open source, GUI-based applications that implement full image analysis pipelines

A full pipeline for applying image-based ML models to image analysis tasks in a versatile way requires software to carry out tasks ranging from image labeling and curation to visualizing results of ML model predictions. In other fields that are actively using image-base ML methods, for example in neuroscience and quantitative behavior, packages have been developed that perform all of these steps within a single piece of software (Mathis *et al*. 2020). Although some efforts are underway to produce similar “all-in-one” packages for analyzing imagery from marine environments (e.g., the VIAME project; Richards *et al*. 2019), and several application-specific packages are already in use (e.g. CoralNet, Lozada-Misa *et al*. 2017; ReefCloud, ReefCloud 2021), most research groups that apply image-based ML models to data from the field still use custom software pipelines that often combine many packages and software modules (see references in Table 3). This is partially driven by the fact that new ML methods are being developed so rapidly that by the time they can be implemented within larger software architectures, they are no longer state-of-the-art. This may simply be a fact of the times, but the difference between using even slightly dated ML methods and not using ML methods at all is much greater than the difference between slightly dated methods and the state-of-the-art. Thus, we believe that creating software architectures that allow users to easily build their own annotated image libraries and to quickly test and evaluate performance of a suite of widely used ML methods may be the single biggest step that can be taken to encourage broader adoption of these methods in marine science.

The transition from largely manual analysis of imagery to ML-based automated analysis is already taking place in other fields, and the availability of free, GUI-based, and actively maintained software packages that integrate all steps in the pipeline has played a major part in facilitating this transition. We point to the *DeepLabCut* package (Mathis *et al*. 2020, Mathis and Mathis 2020, https://github.com/DeepLabCut/DeepLabCut) developed for the study of neuroscience and quantitative behavior from laboratory videos as a potent example of how free, user-friendly, community-supported software can vastly increase the degree to which image-based ML methods are used by a research community, and the utility of such methods for researchers.

The evolving research needs of the marine science community will undoubtedly lead to new priorities, and we do not intend these suggestions to be an exhaustive set of recommendations. However, we believe these steps would go a long way toward making image-based machine learning easier to use, more reliable, and more powerful. As we move toward these goals, it will be crucial to create an open dialogue between researchers who are developing and testing image-based ML methods and researchers who are collecting and working with imagery from the field. The objective of this dialogue should be, in part, to increase fluency on both sides. But such a dialogue will also help fuel the development of novel methods that empower marine scientists to use machine learning to study the ocean in ways that were never before possible.

## Acknowledgements

We thank B. Schlining, B. Martin, and M. Gil for useful discussions, advice, and technical support that improved this manuscript. This work was supported by National Science Foundation grant IOS-1855956 and REU supplement to AMH.

